# Camera-based monitoring of Bogong moths in Alpine Australia reveals drivers of migratory behaviour

**DOI:** 10.1101/2022.05.19.492631

**Authors:** Jesse R A Wallace, Therese Reber, Lana Khaldy, Benjamin Mathews-Hunter, Ken Green, David Dreyer, Eric J Warrant

**Affiliations:** Research School of Biology, Australian National University; Lund Vision Group, Department of Biology, Lund University, Sweden; School of Life and Environmental Sciences, The University of Sydney, Australia; College of Asia and the Pacific, Australian National University

**Keywords:** Bogong moth, camfi, insect conservation, insect flight behaviour, insect monitoring, object detection, wildlife camera, wingbeat frequency

## Abstract

The Bogong moth *Agrotis infusa* is well known for its remarkable annual round-trip migration from its breeding grounds across eastern Australia to its aestivation sites in the Australian Alps, to which it provides an important annual influx of nutrients. Over recent years, we have benefited from a growing understanding of the navigational abilities of the Bogong moth. Meanwhile, the population of Bogong moths has been shrinking. Recently, the ecologically and culturally important Bogong moth was listed as endangered by the IUCN Red List, and the establishment of a program for long-term monitoring of its population has been identified as critical for its conservation. Here, we present the results of two years of monitoring of the Bogong moth population in the Australian Alps using a recently developed method for automated monitoring of flying insects, named Camfi. We found that the evening flights of Bogong moths occur throughout summer, and are modulated by daily weather factors. We present a simple heuristic model of the arrival to and departure from aestivation sites by Bogong moths, and confirm results obtained from fox-scat surveys which found that aestivating Bogong moths occupy higher elevations as the summer progresses. We also present the first recorded observations of the impact of bushfire smoke on aestivating Bogong moths. We observed a dramatic reduction in the size of a cluster of aestivating Bogong moths during the fire, and evidence of a large departure from the fire-affected area the day after the fire. Our results highlight the challenges of monitoring Bogong moths in the wild, and support the continued use of automated camera-based methods for that purpose.

## 1 Introduction

The Bogong moth *Agrotis infusa* is well known for its remarkable annual round-trip migration from its breeding grounds across eastern Australia to its aestivation sites throughout the high mountain areas of New South Wales, Victoria, and the Australian Capital Territory, where it forms aggregations numbering in the millions (reviewed by Warrant et al., 2016). Bogong moth aestivation was first reported during the 19^th^ century (Bennett, 1834; Scott, 1873), but the moths have been known by Aboriginal people in the areas surrounding the Australian Alps for millennia (Keaney et al., 2016; Stephenson et al., 2020). Aboriginal people once converged on these mountainous regions during the spring-summer months to hunt and feast upon the abundant Bogong moth assemblages (Flood, 1996, 1980). In spite of this, Bogong moth migration was not understood until the 1950s, following the thorough studies of Common (1954, 1952).

In recent years, increasing efforts have been made to understand the migration of the Bogong moth from a neuroethological perspective (e.g. Adden et al., 2020; Dreyer et al., 2018; Vries et al., 2017; Warrant et al., 2016), particularly with respect to how Bogong moths navigate. However, an open question remains as to what the proximate triggers for Bogong moth migration are (Warrant et al., 2016). As well as being interesting in its own right, the answer to this question is rapidly becoming critical to the conservation of the unique Australian Alpine ecosystem, which accommodates many species that rely on the annual influx of nutrients brought by the Bogong moth migration (Gibson et al., 2018; Green, 2011, 2003). Concerningly, an estimated 200-fold reduction in the Bogong moth population was observed between the 2016–2017 and 2017–2018 summers, following a slow, but consistent decline since the early 1980s (Green et al., 2021; Mansergh et al., 2019). This has led to the recent listing of the Bogong moth as endangered on the IUCN Red List (Warrant et al., 2021).

The question of what proximate cues trigger Bogong moth migration is complex, and is unlikely to be solved by a single study. Behavioural experiments are laborious, and indeed, to our knowledge, a behavioural paradigm to measure the timing of a Bogong moth’s migration in response to controlled stimuli has yet to be developed. In the meantime it therefore seems prudent to make quantitative measurements of Bogong moth migratory timing in the wild. This will at least enable us to determine what proximal factors are correlated with the behaviour, which will greatly assist in narrowing the search-space for future experimentation.

Useful progress to this end has been made through long-term monitoring of migrating insects using vertical radar deployed on the Bogong moth migratory route (e.g. Hao et al., 2020). However, reliable monitoring of Bogong moths in their breeding grounds remains an unsolved challenge (Wintle et al., 2021). At the end of their spring migration, a number of methods have been used to monitor Bogong moths close to their aestivation sites, including light trapping (Gibson et al., 2018; Wintle et al., 2021), light beam surveys, (Monk, 2021), aestivation site surveys (Caley and Welvaert, 2018; Green et al., 2021), ski surveys of Bogong moth carcasses on the snow (Green et al., 2021), and fox scat surveys (Green, 2010; Green and Osborne, 1981). Each of these methods have their idiosyncrasies, and are to varying degrees laborious, limiting their utility for large-scale long-term monitoring programs, such as the 100-site Bogong moth monitoring program recommended by Wintle et al. (2021).

In this paper, we present the results of two years of monitoring of Bogong moth flight activity near aestivation sites in the Australian Alps using wildlife cameras, and a newly developed method described by Wallace et al. (2021). We show that by monitoring the sites for the full span of the Bogong moth aestivation season, we are able to infer the arrival and departure dates of the moths from those sites. Moreover, we are able to quantitatively analyse the evening twilight flight activity of the aestivating Bogong moths described by Common (1952) over the entire duration of the summer, providing strong preliminary evidence for weather being an important driver of flight behaviour—and by extension, migratory behaviour—in the moth. Our results, and indeed the method we have developed to obtain them, may be of interest to land managers and conservationists who seek to measure the ongoing effects of management practices on the Bogong moth population, and to monitor it more generally.

## 2 Methods

The methods employed in this study closely follow those described by Wallace et al. (2021), and are described briefly below. Weather data were obtained from weather stations close to the camera sites (Bureau of Meteorology, 2021).

### 2.1 Camera placement and settings

Study sites were selected for their proximity to known Bogong moth aestivation sites. In the first study season (2019–2020), cameras were placed outside a Bogong moth aestivation site in a boulder field near the summit of Mt Kosciuszko, NSW, and outside two aestivation sites near the formerly unnamed peak now known as Ken Green Bogong (referred to in this paper as K.G. Bogong), near South Rams Head, NSW. In the second study season (2020–2021), a site near the summit of Mt. Gingera, ACT/NSW, which has been subject to a number of previous studies (Caley and Welvaert, 2018; Common, 1954; Keaney et al., 2016) was added. Data obtained in November 2019 from a boulder field near Cabramurra, NSW, by Wallace et al. (2021) were also included for certain analyses.

During the 2019–2020 season, a single camera was also placed inside the aestivation cave on K.G. Bogong, facing towards a cluster of aestivating Bogong moths (referred to as the “observation cluster”). This camera was not used for automated annotation, although occasionally flying moths were seen inside the cave. By the end of the season this camera had been flooded and was no longer usable.

### 2.2 Image annotation

A total of 109912 images of the sky was obtained during this study. Of these, we manually annotated 33780 images for Bogong moths. Of the 33780 manually annotated images, 4223 contained at least one annotation. We kept 200 of the 4223 images as a test set, combining them with the test set used by Wallace et al. (2021). The remaining 4023 of these images were combined with the training set used by Wallace et al. (2021), for a total training set of 4901 images. The Camfi annotation model was retrained on this image set and the newly trained model was evaluated on the combined test set. The newly trained model has been included with a recent release of Camfi, which is available at https://github.com/J-Wall/camfi.

All images obtained in this study were then automatically annotated using Camfi with the newly trained model. Wingbeat frequencies of each annotation were then measured using Camfi. For further analyses, the automatically obtained annotations were filtered by prediction score, wingbeat SNR, and wingbeat frequency. In particular, annotations with prediction scores less than 0.8, wingbeat SNR outside of the range [1, 50], or wingbeat frequency outside of the range [27, 78] Hz were excluded.

### 2.3 Data analysis

Of primary interest is the relative daily abundance of flying Bogong moths at the study sites and across the study period. We measured this by counting moth detections occurring during evening twilight, noting the number of images collected at a given site during the evening twilight of a given day as the exposure variable.

Various daily abiotic factors were regressed against counts of Bogong moths detected by Camfi during evening twilight. Before performing the regression, the Pearson correlations between each pair of factors were calculated, and highly correlated factors were removed using a greedy recursive algorithm. The algorithm proceeded by selecting the most highly correlated pair of factors, then removing the factor in the pair which was less well correlated with the evening detection count. The algorithm terminates when no pair of factors had a Pearson *R*^2^ greater than a specified threshold, which we set to 0.3. The remaining factors were then jointly regressed against the evening detection count using a Poisson regression with image count as the exposure variable.

## 3 Results

An evaluation of the performance of the newly trained annotation model is presented in Appendix A.1. Overall, every evaluation metric marginally improved with respect to the previous Camfi annotation model (Wallace et al., 2021), with the exception of average precision, which slightly worsened. This is presumably a consequence of the present dataset containing images taken in a wider variety of lighting conditions.

Strong peaks in activity were observed during evening twilight across all study sites (Fig. 1, *right panel*), although pronounced peaks were not seen during morning twilight (Fig. 1, *left panel*). Evening twilight detection counts were highly variable, but clearly show that Bogong moths departed from the lower elevation sites (Mt. Gingera and K.G. Bogong) earlier in the season than from higher elevation sites, i.e. Mt. Kosciuszko (Fig. 2 for the 2019–2020 summer and Fig. 3 for the 2020–2021 summer). The camera placed at the K.G. Bogong site fell from its mount towards the end of the season (Fig. 3, *lower panel, shaded region*), reducing the camera’s view of the sky by about half, however it appears that the majority of moths had already left the area by the time this happened.

**Figure 1.**
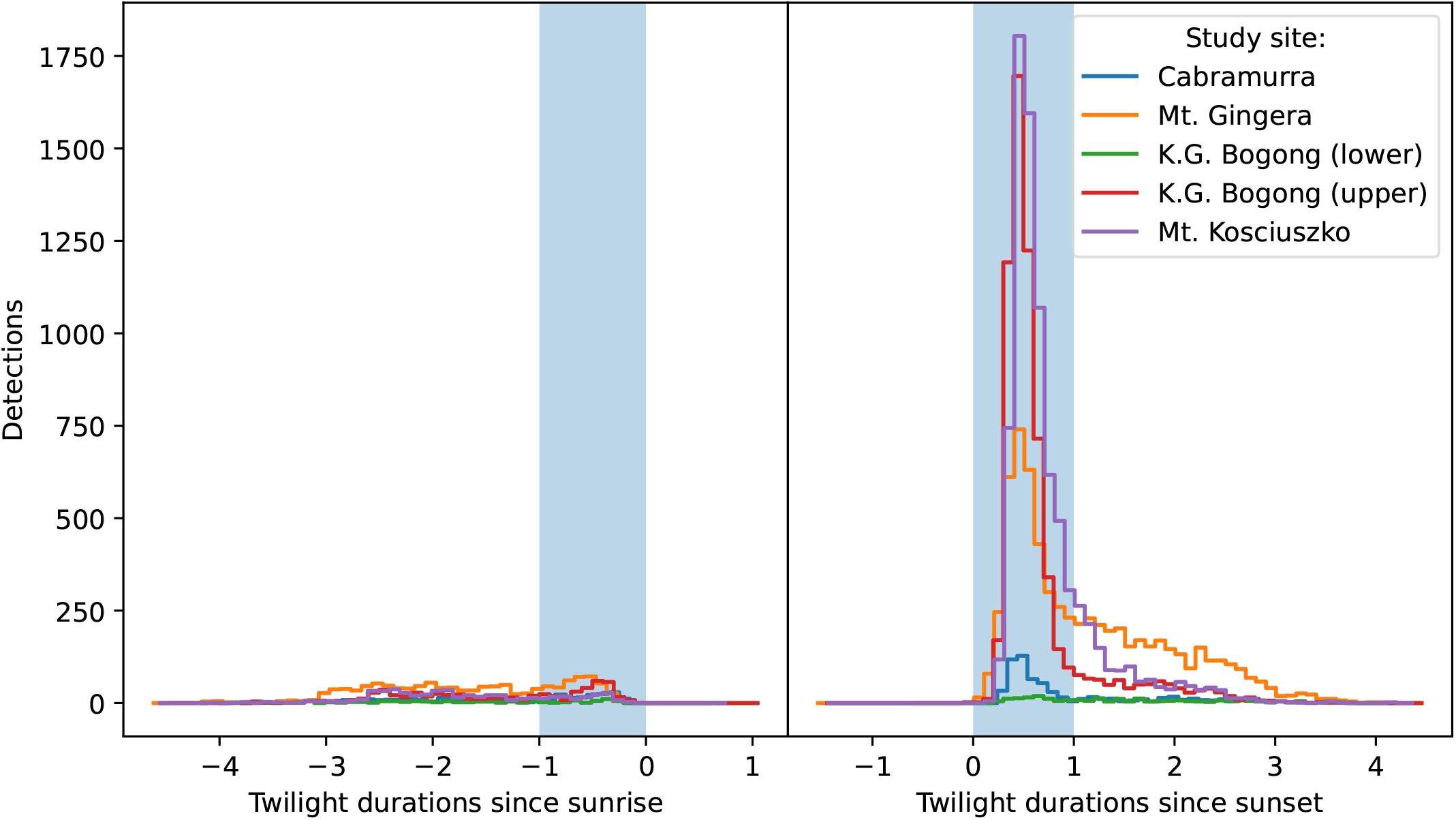
Total number of moth observations by time relative to sunrise (*left*, scaled by the duration of morning twilight) does not show peak in activity during morning twilight (*blue shaded region*), with the slight exception of K.G. Bogong (upper) and Mt Gingera sites, which show small peaks. Total number of moth observations by time relative to sunset (*right*, scaled by the duration of evening twilight) shows peak in activity during evening twilight (*blue shaded region*) across all study sites. Data from Cabramurra boulder field site is from Wallace et al. (2021).

**Figure 2.**
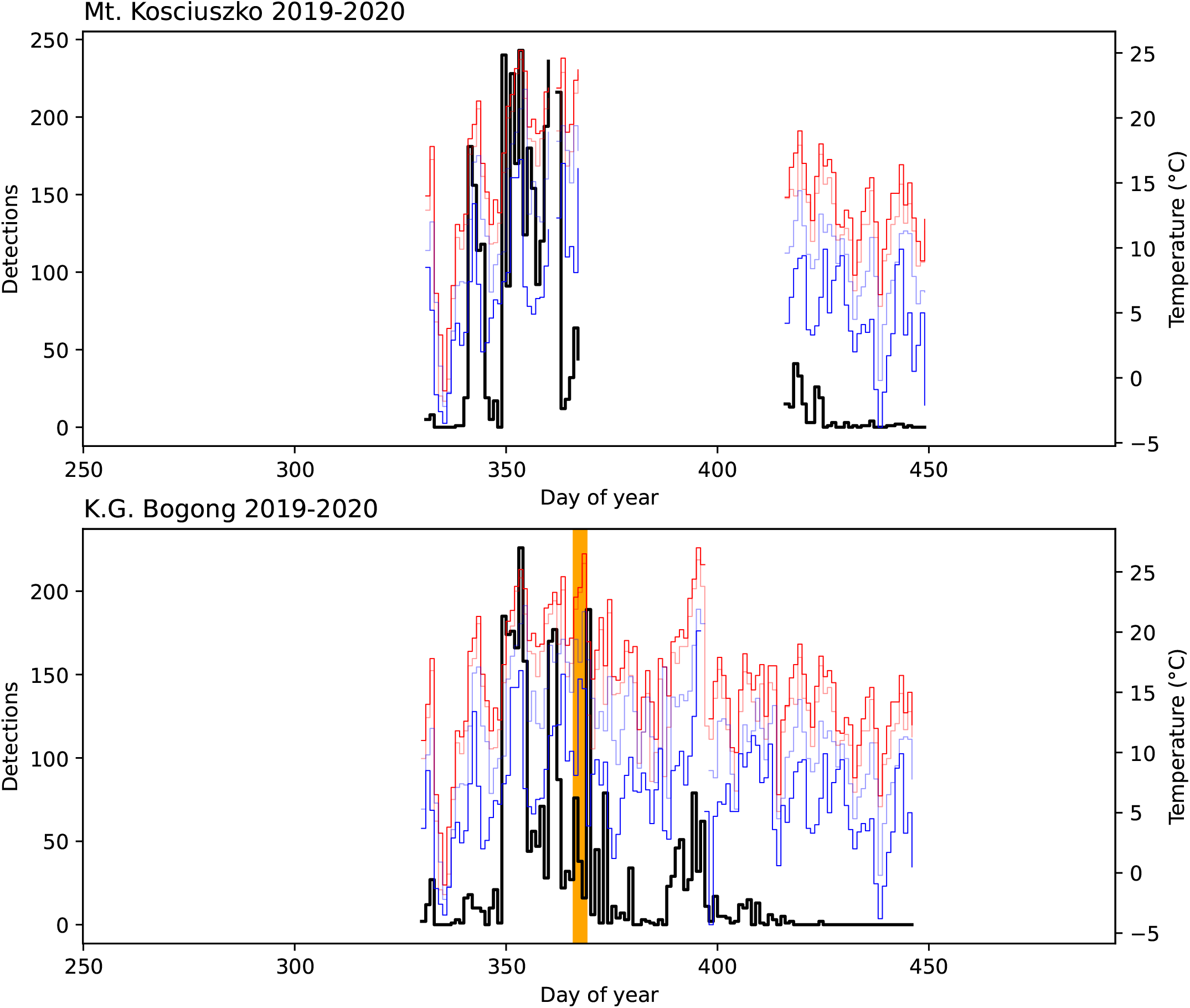
Number of Bogong moth detections (*black*) for each study day in the 2019–2020 summer season outside Bogong moth aestivation sites on Mt. Kosciuszko and K.G. Bogong, NSW, shown with daily temperatures recorded at Thredbo Top Station (Bureau of Meteorology, 2020): maximum (*red*), minimum (*blue*), 9 am (*light blue*), and 3 pm (*light red*). Data are missing for a portion of the season at the Mt. Kosciuszko site due to a camera malfunction. *Orange span* indicates a bushfire event which occurred 1 km SW of K.G. Bogong (the fire did not reach the site, although there was high levels of smoke in the air which would have entered the site).

**Figure 3.**
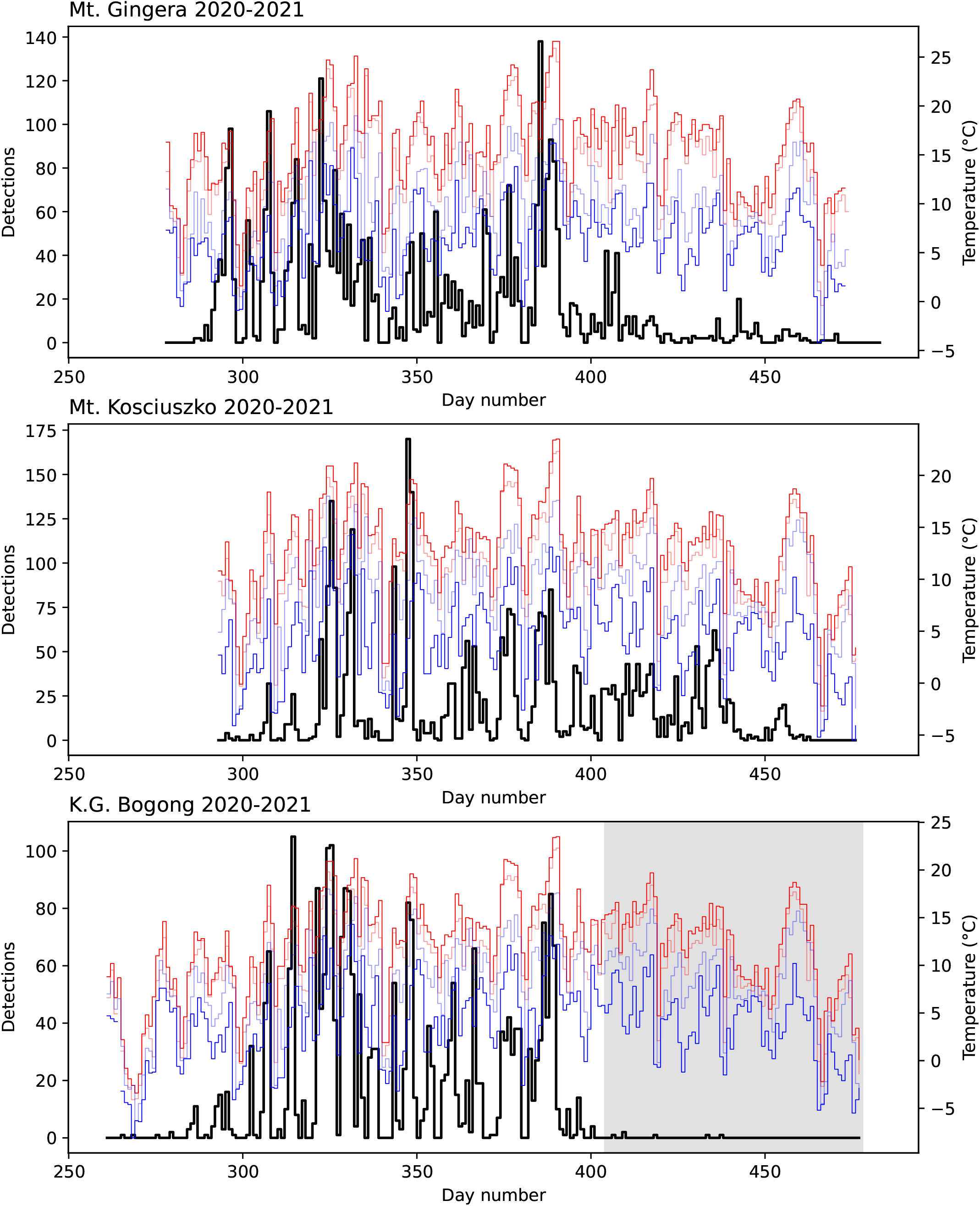
Number of Bogong moth detections (*black*) for each study day in the 2020–2021 summer season outside Bogong moth aestivation sites on Mt. Gingera, Mt. Kosciuszko, and K.G. Bogong, NSW, shown with daily temperatures recorded at Mt Ginini and Thredbo Top Station (Bureau of Meteorology, 2021): maximum (*red*), minimum (*blue*), 9 am (*light blue*), and 3 pm (*light red*). *Shaded region* on lower plot indicates period where camera had fallen from its mount, reducing its view of the sky by about half.

### 3.1 Predictors of activity

A total of ten abiotic factors were found to be significantly correlated with evening twilight counts of flying Bogong moths (Fig. 4a). A greater number of moths were observed at higher elevation sites (Fig. 4b) and when twilight duration was longer (i.e. in the middle of summer, Fig. 4d). Study year was also positively correlated with moth counts, suggesting that the Bogong moths were more abundant in the 2020–2021 summer than in the 2019–2020 summer. The most important weather factors were daily maximum temperature (which had a positive effect on moth counts, Fig. 4c) and maximum wind speed (which had a negative effect on moth counts, Fig. 4e). Note that very few Bogong moths were observed flying on days which had maximum temperatures lower than 10°C (Fig. 4c). Daily temperature range, relative humidity (measured at 9 am), and daily minimum temperature were negatively correlated with moth counts, while latitude and rainfall were positively correlated with moth counts. Scatter plots of all covariates in our model are shown in Fig. A.2, and residuals of the fitted model are shown in Fig. A.3 (both in Appendix A.2).

**Figure 4.**
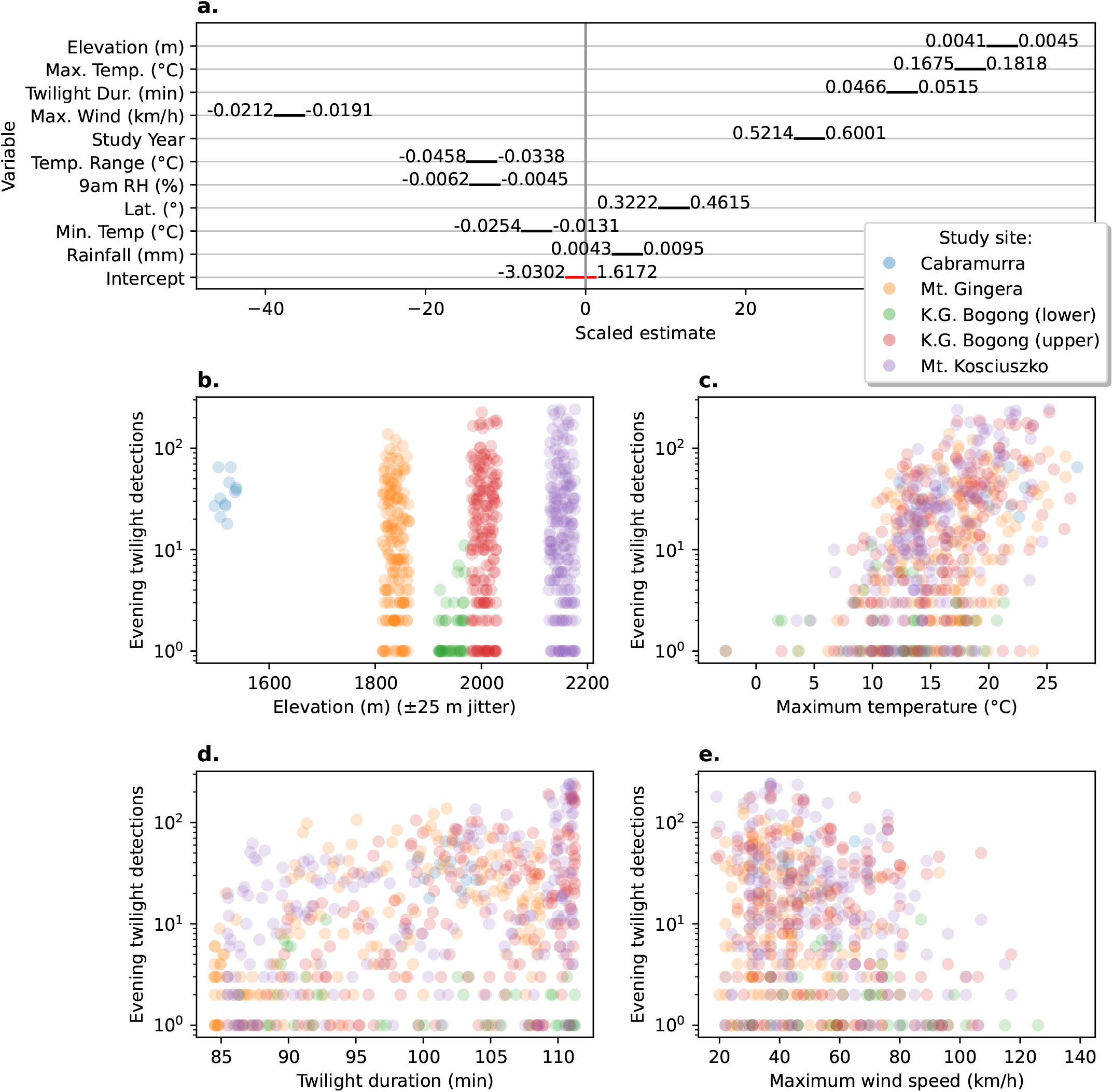
Effect-sizes and plots of number of detections during evening twilight against significantly associated abiotic factors. **a**. Scaled estimates of effect size of abiotic factors on Bogong moth evening flight intensity (as measured by number of Camfi detections) from a mixed-effect Poisson generalised linear model of detections against these factors. *Black bars* show 95% confidence interval of estimates (scaled by effect size). Values to either side of *black bars* represent bounds of 95% confidence interval in the units of the respective factor (corresponding to the gradient of the regression, in that dimension). Negative values indicated that increases in the value of the factor lead to a decrease in moth counts (and positive values, the opposite). **b**. Scatter plot of detections per evening twilight by elevation. Random fluctuation (jitter) is applied to elevation to increase readability. **c**. Scatter plot of detections per evening twilight by daily maximum temperature. **d**. Scatter plot of detections per evening twilight by duration of evening twilight. **e**. Scatter plot of detections per evening twilight by daily maximum wind gust speed. Points in *b–e* are coloured by study site, as per study-site key (*right, towards top*).

### 3.2 Arrival and departure of Bogong moths

During the 2020–2021 summer season, the cameras were placed at the aestivation sites before the Bogong moths had arrived, and removed after they had left. This means the detection data obtained from those cameras contain information regarding the arrival and departure dates of the Bogong moths. For example, at Mt. Gingera, the first Bogong moth was detected on the 13^th^ October (Fig. 3, *top panel, day 286*). At K.G. Bogong, this date was 11^th^ October (Fig. 3, *bottom panel, day 284*). A few detections were made prior to this date, however upon inspection these were found to be false-positives caused by rain. The first detection of a Bogong moth at Mt. Kosciuszko was on 22^nd^ October (Fig. 3, *middle panel, day 295*) although cameras were only placed there on 20^th^ October, so it is possible that some moths had arrived earlier. However, snow was still present on the ground in front of the aestivation site on Mt. Kosciuszko until it melted on 23^rd^ October (JRAW, personal observation), so if there was an earlier arrival, it was probably only by a few days.

While records of earliest arrival are interesting, they are also subject to substantial noise, owing to the fact that the marginal probability of a particular moth being detected at all is very small. Earliest arrivals are also not necessarily representative of the predominant behaviour in the population. Therefore, we would like to use the detection data across the entire season to model the arrival and departure of the majority of the population. To do this, we propose a simple heuristic model of the evolving relative abundance of evening-flying Bogong moths in an area, as the moths arrive at—and later depart from—the area (Fig. 5). This model is based purely on detection data, and is independent of weather factors, etc.

**Figure 5.**
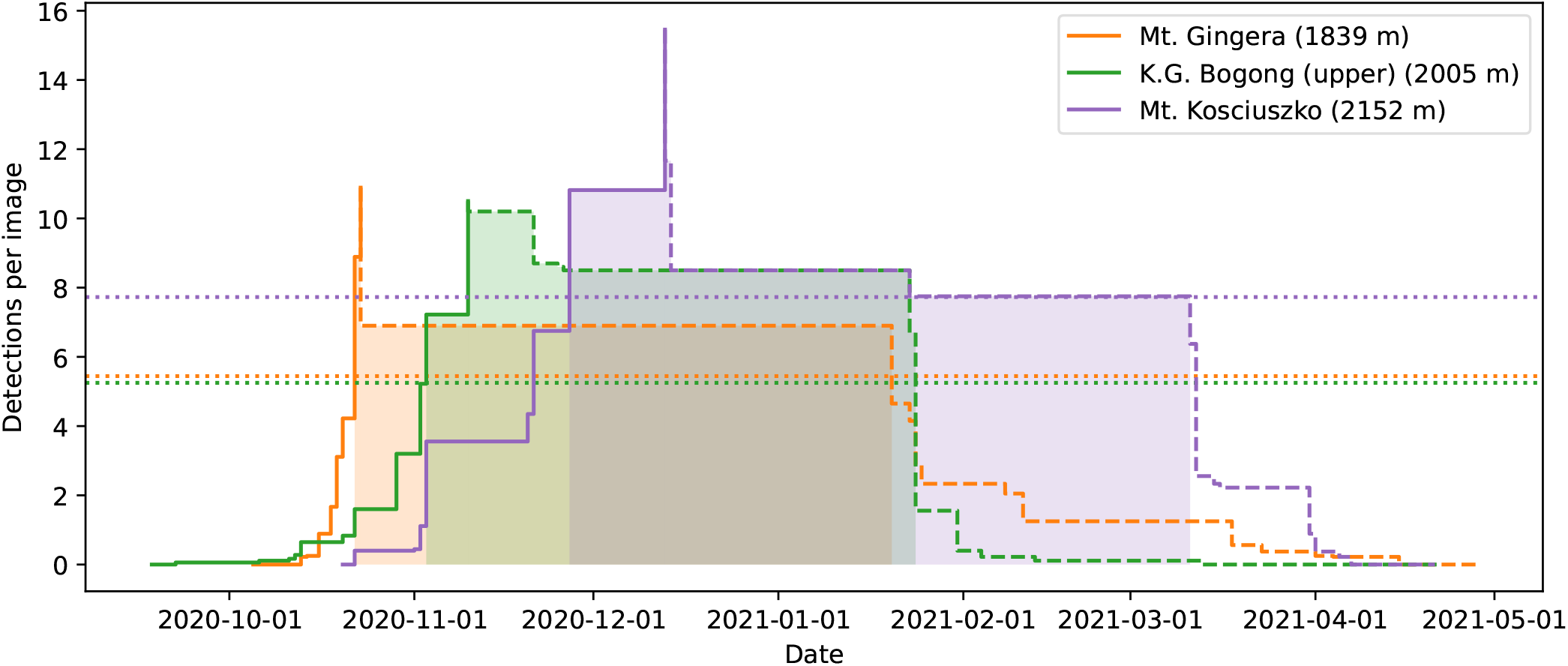
A simple heuristic model of the arrival and departure of Bogong moths to summer aestivation sites applied to data obtained from automated camera monitoring in the summer of 2020–2021. *Solid lines:* Cumulative maximum detections per image, plotted until the date that absolute maximum is reached, for the respective location. This roughly models the arrival of moths to the location. *Dashed lines:* Reverse-cumulative maximum of detections per image, plotted from date of absolute maximum, for the respective location. This roughly models the sum of departure and mortality of moths from the location. *Dotted lines:* Show half of the maximum detections per image for respective location (for calculating median date of arrival and departure). Dates after median date of arrival, and before median date of departure for each location are *shaded*. Elevations shown are of the camera placement, rather than the summit elevations of the mountains.

The model separates the aestivation of Bogong moths at a particular site into two phases; arrival and departure. In the arrival phase (Fig. 5, *solid lines*), the relative abundance of aestivating moths is modelled by the cumulative maximum of the mean number of detections per image over all preceding evenings. The arrival phase ends on the day where this value reaches its maximum across the entire summer. The departure phase (Fig. 5, *dashed lines*) is modelled similarly, this time using the reverse-cumulative maximum.

An obvious set of descriptive statistics arise; namely, median date of arrival and median date of departure. These are shown for each study site in the 2020–2021 season in Fig. 5 (*shaded areas*) and in Table 1. A clear signal of Bogong moths arriving at higher elevation aestivation sites later than lower elevation sites is present (Fig. 5, *solid lines*; Table 1). This trend appears to also apply to departures, albeit slightly less clearly (Fig. 5, *dashed lines*; Table 1).

**Table 1.** Median date of arrival (*A*_1/2_) and departure (*D*_1/2_) of Bogong moths from aestivation sites during 2020–2021 summer. Elevations shown are of the camera placement, rather than the summit elevations of the mountains. 3-day average maximum is calculated across the 3 days preceding the date listed (inclusive), from the nearest weather station (Bureau of Meteorology, 2020) assuming an adiabatic lapse rate of 9.1°C/1000 m elevation (Green, 2014).

| Date | Mt. Gingera (1839 m) | K.G. Bogong (2005 m) | Mt. Kosciuszko (2152 m) |
| --- | --- | --- | --- |
| Mt. Gingera $A_{1/2}$<br>(2020-10-22) | 14.0°C (std = 2.0°C) | 11.2°C (std = 1.7°C) | 9.8°C (std = 1.7°C) |
| K.G. Bogong $A_{1/2}$<br>(2020-11-03) | 14.0°C (std = 2.5°C) | 14.0°C (std = 4.1°C) | 12.7°C (std = 4.1°C) |
| Mt. Kosciuszko $A_{1/2}$<br>(2020-11-27) | 20.1°C (std = 2.4°C) | 17.3°C (std = 1.9°C) | 16.0°C (std = 1.9°C) |
| Mt. Gingera $D_{1/2}$<br>(2021-01-20) | 18.4°C (std = 2.2°C) | 16.1°C (std = 0.9°C) | 14.7°C (std = 0.9°C) |
| K.G. Bogong $D_{1/2}$<br>(2021-01-24) | 24.3°C (std = 1.6°C) | 20.7°C (std = 2.3°C) | 19.4°C (std = 2.3°C) |
| Mt. Kosciuszko $D_{1/2}$<br>(2021-03-11) | 16.4°C (std = 1.4°C) | 14.4°C (std = 1.5°C) | 13.0°C (std = 1.5°C) |

To assess whether temperature could explain the later arrival and departure of Bogong moths at higher elevations, we calculated the 3-day average maximum temperature at each location in the lead-up to the median arrival and departure (Table 1). Notably, temperatures were relatively high at lower elevation sites (Mt. Gingera: 20.1°C, K.G. Bogong: 17.3°C) in the days leading up to the median *arrival* date at the higher-elevation Mt. Kosciuszko site (2020-11-27; Table 1). Also, temperatures in the lead-up to median *departure* dates at the lower elevation sites were relatively high, respectively (Mt. Gingera: 18.4°C, K.G. Bogong: 20.7°C), while pre-departure temperatures at Mt. Kosciuszko were comparatively cool (13.0°C; Table 1).

### 3.3 Impact of January 2020 bushfire

On 4^th^ January 2020, a major bushfire which had been burning in the area in the preceding few days (Fig. 2, *orange span*) came within 1 km of the K.G. Bogong site. Despite the thick smoke, Bogong moths were seen flying outside their aestivation cave (Appendix A.3 Fig. A.4). The following day, a large number of flying Bogong moths were detected, presumably indicating a departure of a portion of the moths from the site (Fig. 2, peak to the right of *orange span*).

A reduction in the number of aestivating Bogong moths on K.G. Bogong during the bushfire was reflected by our observation cluster, which dramatically reduced in size over the course of the fire (Fig. 6). Notably, a significant portion of this reduction happened during the day (Fig. 6b–c), despite Bogong moths typically being night-active. The remaining cluster (Fig. 6c) did not change much during the following few weeks before the camera was flooded on 20^th^ January.

**Figure 6.**
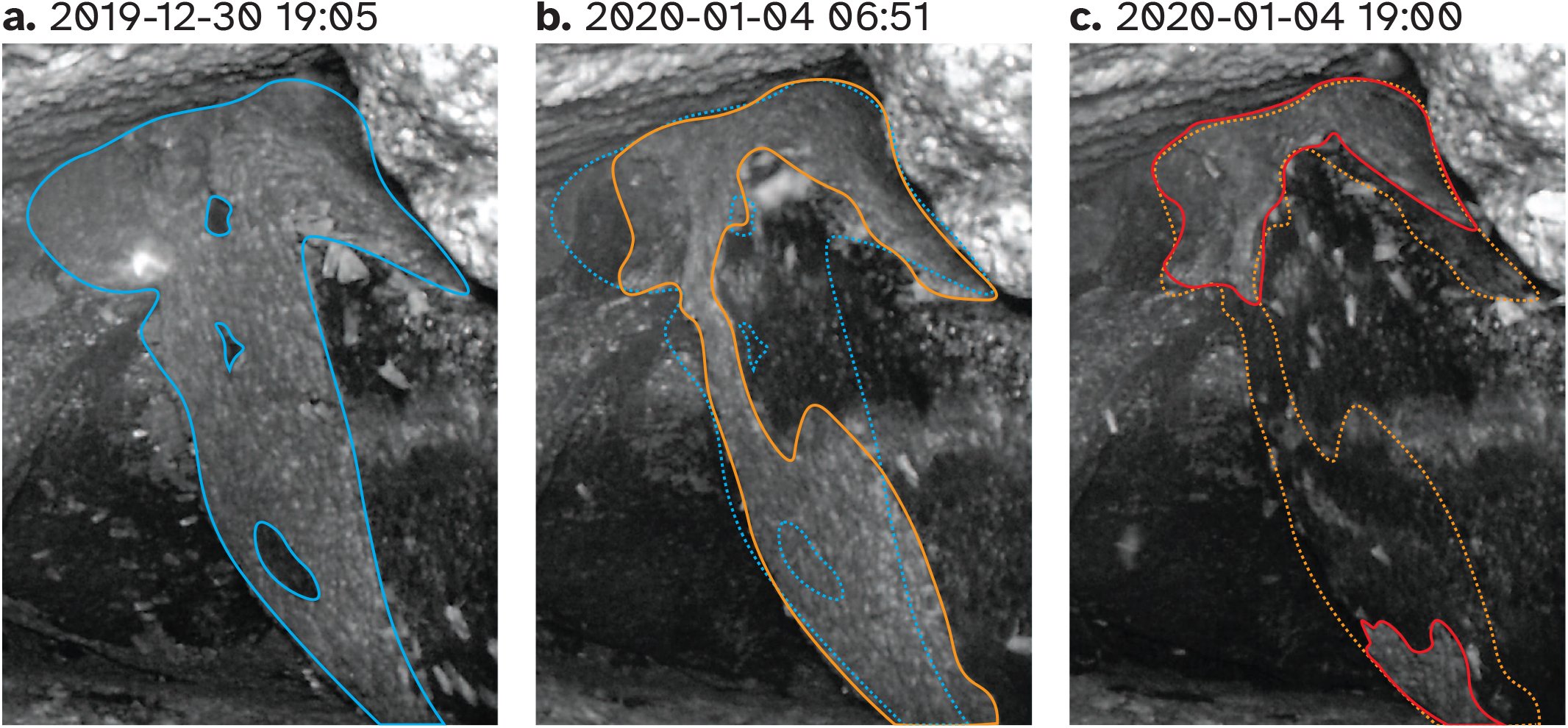
Progression of cluster of aestivating Bogong moths in cave on K.G. Bogong over the worst few days for Kosciuszko National Park during the 2019–2020 bushfire season. **a**. Prior to bushfire. Cluster is outlined with *solid blue trace*. **b**. Morning of 4^th^ January 2020, the day which saw the bushfire come within 1 km of the site. Cluster of Bogong moths is outlined with *solid orange trace*. Trace from *a* is overlaid for comparison (*dotted blue trace*). **c**. Evening of 4^th^ January 2020. Cluster of Bogong moths is outlined with *solid red trace*. Trace from *b* is overlaid for comparison (*dotted orange trace*). Times shown are Australian Eastern Daylight Time (AEDT; UTC+11:00).

## 4 Discussion

Our results clearly and quantitatively demonstrate that the summer flights of Bogong moths described by Common (1954) occur predominantly during evening twilight, and occur throughout the Bogong moth’s entire summer aestivation. Furthermore, the intensity of these flights is modulated by daily weather factors, with Bogong moths favouring warmer evenings with lower wind speeds for flying, confirming that the patterns seen by Wallace et al. (2021) hold over the entire summer.

For simplicity’s sake, we chose to use a relatively simple linear model relating moth counts to various abiotic factors. If long-term monitoring of Bogong moths continues using our camera-based method, and an increasing number of years of data become available, more complex models will become more appropriate. For example, we have modelled our study year as having a linear effect on moth counts. For a two-year study, this is valid, however when additional years are added, this should be changed to a random effect (or perhaps an effect depending on annual climactic factors, depending on the research question).

Our survey of three known Bogong moth aestivation sites over the summer of 2020–2021 shows that occupation of higher elevation aestivation sites by Bogong moths occurs later in the season than lower elevation sites. There are three possible explanations for the later arrival dates at the higher elevations: 1) The higher sites are blocked by snow (or are otherwise unsuitable) earlier in the season, but are open later to allow occupation by later arrivals of Bogong moths coming directly from the breeding grounds, 2) Bogong moths from lower sites move higher as summer progresses, or 3) a combination of 1 and 2.

Blockage of high-elevation aestivation sites by snow (possibility 1) would certainly prevent Bogong moths from occupying those sites, but this does not appear to be a satisfactory explanation for the delay we observed, as most of the remaining snow near the highest site (Mt. Kosciuszko) melted on 23^rd^ October 2020, more than an entire month prior to the median arrival of Bogong moths at this location. Additionally, sub-zero temperatures have been recorded in occupied Bogong moth aestivation sites (Green et al., 2021), so it seems unlikely that low temperatures alone would have *prevented* Bogong moths from migrating directly to high-altitude sites early in the season.

On the other hand, high temperatures do appear to be a reasonable explanation for Bogong moths *avoiding* lower elevation sites, which could motivate movement to higher elevations (possibility 2 or possibility 3). The median arrival date at the highest elevation site (Mt. Kosciuszko) coincided with lower elevation sites experiencing 3-day average maximum temperatures above 16°C (Table 1). Incidentally, Green et al. (2021) estimated that 16°C is the maximum temperature (inside a cave) that permits aestivation. Interestingly, 3-day average maximum temperatures leading up to the median *departure* dates at Mt. Gingera and K.G. Bogong were 18.4°C and 20.7°C at those locations, respectively (i.e. well above 16°C; Table 1). However, the 3-day average maximum temperature at Mt. Kosciuszko leading up to its median *departure* date was lower, at just 13.0°C. Therefore, it could be that departures from the Mt. Gingera and K.G. Bogong sites were motivated by high temperatures (and resulted in movements to higher elevations), while departures from Mt. Kosciuszko were motivated by temperatures falling (to 13°C), indicating the approaching autumn, thus triggering the return migration to the breeding grounds.

Notably, possibility 2 is also supported by previous results from fox scat surveys (Green, 2010), which showed a departure of Bogong moths from sub-alpine areas into alpine areas as the summer progressed. However, properly disentangling each of these possibilities requires observations of the movements (or lack thereof) of Bogong moths between elevations during the summer, after their arrival in the mountains. Such observations could be made using a similar method to that used in this study, with cameras deployed in elevation transects on a single mountain.

Our simple model of Bogong moth arrival and departure is robust to periods of evening-flight inactivity (e.g. due to unfavourable weather conditions for flight), and follows naturally from the following two heuristics: 1) the maximum relative density of flying moths in the vicinity of an aestivation site is representative of the relative abundance of aestivating moths in the area, and 2) most Bogong moths arrive at and leave from the vicinity of an aestivation site on relatively few nights (so evening flights with lower relative density are generally station-keeping movements, rather than migrations to other aestivation sites or returning to the breeding grounds). As with the model for relating evening flights to abiotic factors, the arrival and departure model could perhaps be extended to include—depending on the research question—the effects of other factors, such as daily weather and annual climate.

Fire appears to have a pronounced effect on assemblages of aestivating Bogong moths. We observed a marked reduction in the size of our observation cluster of aestivating Bogong moths during the day that a bushfire came close to the site (but did not affect it directly). Presumably this was mediated by smoke entering the cave and disturbing the moths. Interestingly, from flight data, we observed (what we assume to be) a large departure of Bogong moths from a bushfire-affected area the day *after* the fire, and thus the day after the marked reduction in the size of the observation cluster. It could be that the reduction of the observation cluster on the day of the fire was caused by Bogong moths falling from their perch on the cave wall, but remaining inside the cave, rather than perishing or departing that day. These moths could then have departed the following day when conditions outside were less dangerous.

In recent years, especially since their dramatic population crash in 2017 (Green et al., 2021; Mansergh et al., 2019), Bogong moths have enjoyed increased attention from those interested in their conservation, and in the conservation of the Australian Alpine ecosystem more generally. In particular, the need for the implementation of a long-term monitoring program for Bogong moths for the ongoing conservation of the Australian Alpine ecosystem has been identified (Wintle et al., 2021).

The high level of variability in counts we observed across each night of our study highlights the importance of regular measurements throughout the summer for such a monitoring program. Not only does the proportion of Bogong moths flying on a given night depend on the weather, but the Bogong moth population also moves between aestivation sites over the course of the summer, complicating the interpretation of sparse data collected from infrequent light-trapping surveys, particularly when these surveys do not simultaneously collect counts from multiple locations.

Ideally, a long-term monitoring program for Bogong moths would collect counts of moths every day at every study site from mid-September until mid-May, completely covering the Bogong moth aestivation season. For a large-scale program with many study sites, light trapping or similarly labour-intensive approaches would be a costly undertaking. Conversely, a comparatively “hands-off” and non-invasive approach (i.e. not involving trapping), such as the automated camera-based monitoring method used in this study, could be relatively easily and inexpensively scaled, without the need for a large team of dedicated light-trappers. For instance, by replacing the wildlife cameras used in this study with permanent solar-powered and Internet-connected camera stations, the data acquisition portion of a large-scale Bogong moth monitoring program could be completely automated, with visits to study sites only needed for placement, maintenance, and eventual retrieval of the equipment. Such a program would produce an incredibly rich and informative dataset for ongoing efforts to model the dynamics of the Bogong moth population, and their migration to and from the Australian Alps.

Finally, one could also imagine a wide variety of possible study designs using the method, as it provides the opportunity to trivially increase the number of cameras at each site, or change the frequency of image captures to address specific research questions surrounding the behaviour of the moths.

## 5 Author contributions

EJW, DD, and JRAW conceived the project. JRAW, TR, BMH, DD, EJW, and KG performed the fieldwork. KG provided essential knowledge of the locations of Bogong moth aestivation sites. TR, LK, BMH, and JRAW performed the manual annotations of images of flying moths. JRAW analysed the data. JRAW wrote the first draft of the manuscript, with input from EJW.

## 6 Acknowledgements

EJW and JRAW are grateful for funding from the European Research Council (Advanced Grant No. 741298 to EJW), and the Royal Physiographic Society of Lund (to JRAW). JRAW is thankful for the support of an Australian Government Research Training Program (RTP) Scholarship. EJW holds Scientific Permits for collection and experimental manipulations of Bogong moths in several alpine national parks and nature reserves (NSW Permit SL100806). We are extremely grateful to Dr. Peter Caley for useful discussions, and to Dr. Stanley Heinze for essential collaboration and providing computational resources.

## A Appendix

### A.1: Automatic annotation evaluation

Automatic annotation performance was evaluated using a test set of 200 images, as well as the full set of 33780 manually annotated images. Evaluation metrics for both sets are presented in Table A.1. Each metric was similar across both image sets, indicating that the annotation model has not suffered from over-fitting. This is also supported by the contour plots of prediction score vs. IoU, polyline Hausdorff distance, and polyline length difference (Fig. A.1b,c,d, respectively). These plots show similar performance on both the full image set (33780 images) and the test set (200 images). Furthermore, they show that prediction scores for matched annotations (automatic annotations which were successfully matched to annotations in the manual ground-truth dataset) tended to be quite high, as did the IoU of those annotations, while both polyline Hausdorff distance *d*_*H*_ and polyline length difference Δ*L* clustered relatively close to zero. The precision-recall curves of the automatic annotator (Fig. A.1e) show similar performance between the image sets, and show a drop in precision for recall values above 0.4, possibly due to poorer performance for images which were taken in “day mode” (without infra-red flash). Training took 17414 iterations and completed in less than 2 h (Fig. A.1a) on a machine with two 8-core Intel Xeon E5-2660 CPUs running at 2.2GHz and a Nvidia T4 GPU.

**Table A.1:** Automatic annotation performance metrics for 2019–2021 study when tested against the full manually-annotated image set (33780 images), and the test set (200 images). Performance metrics calculated are average precision *AP*_50_, mean bounding-box intersection over union 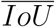, mean Hausdorff distance of polyline annotations 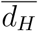, mean signed length difference of polyline annotations 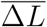, and the standard deviation of signed length difference of polyline annotations *σ*_Δ*L*_. Definitions of these metrics follow those of Wallace et al. (2021). ^†^*AP*_50_ was calculated on the set of images with at least one manual annotation, rather than the full set of 33780 images.

| Image set | $AP_{50}$ | $\overline{IoU}$ | $\overline{d_H}$ | $\overline{\Delta L}$ | $\sigma_{\Delta L}$ |
| --- | --- | --- | --- | --- | --- |
| Full set | 0.426<sup>†</sup> | 0.835 | 25.9 | -2.93 | 39.3 |
| Test set | 0.420 | 0.844 | 21.5 | -0.80 | 28.5 |

**Figure A.1:**
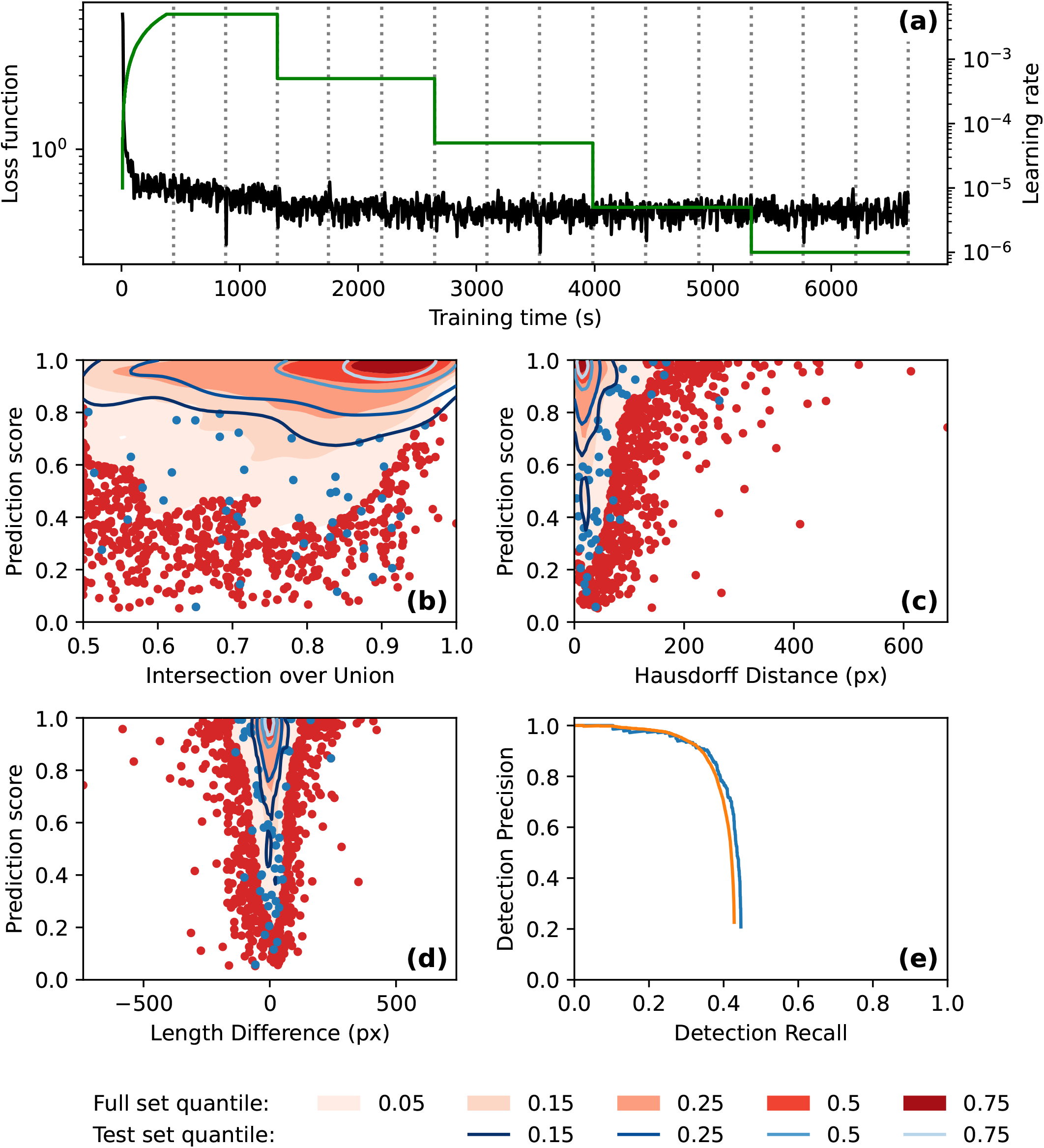
Automatic annotation evaluation plots for 2019–2021 study. **(a)** Automatic annotation model training learning rate schedule (*green*) and loss function (*black*) over the course of training. Epochs (complete training data traversal) are shown with dotted vertical lines. **(b)-(d)** Gaussian kernel density estimate contour plots of prediction score vs. **(b)** bounding box intersection over union,**(c)** polyline Hausdorff distance, and **(d)** polyline length difference, for both image sets. Contours are coloured according to density quantile (key at bottom of figure). “Full set” refers to the set of 33780 images which were manually annotated. “Test set” refers to the set of 200 images with at least one annotation which were not used during model training. In each plot, data which lie outside of the lowest density quantile contour are displayed as points. **(e)** Motion blur detection precision-recall curve, generated by varying prediction score threshold. The precision-recall curve for the test set (200 images) is shown in *blue*, and the precision-recall curve for the set of 4223 images which had at least one manual annotation is shown in *orange*.

### A.2: Bogong evening twilight flight covariates

**Figure A.2:**
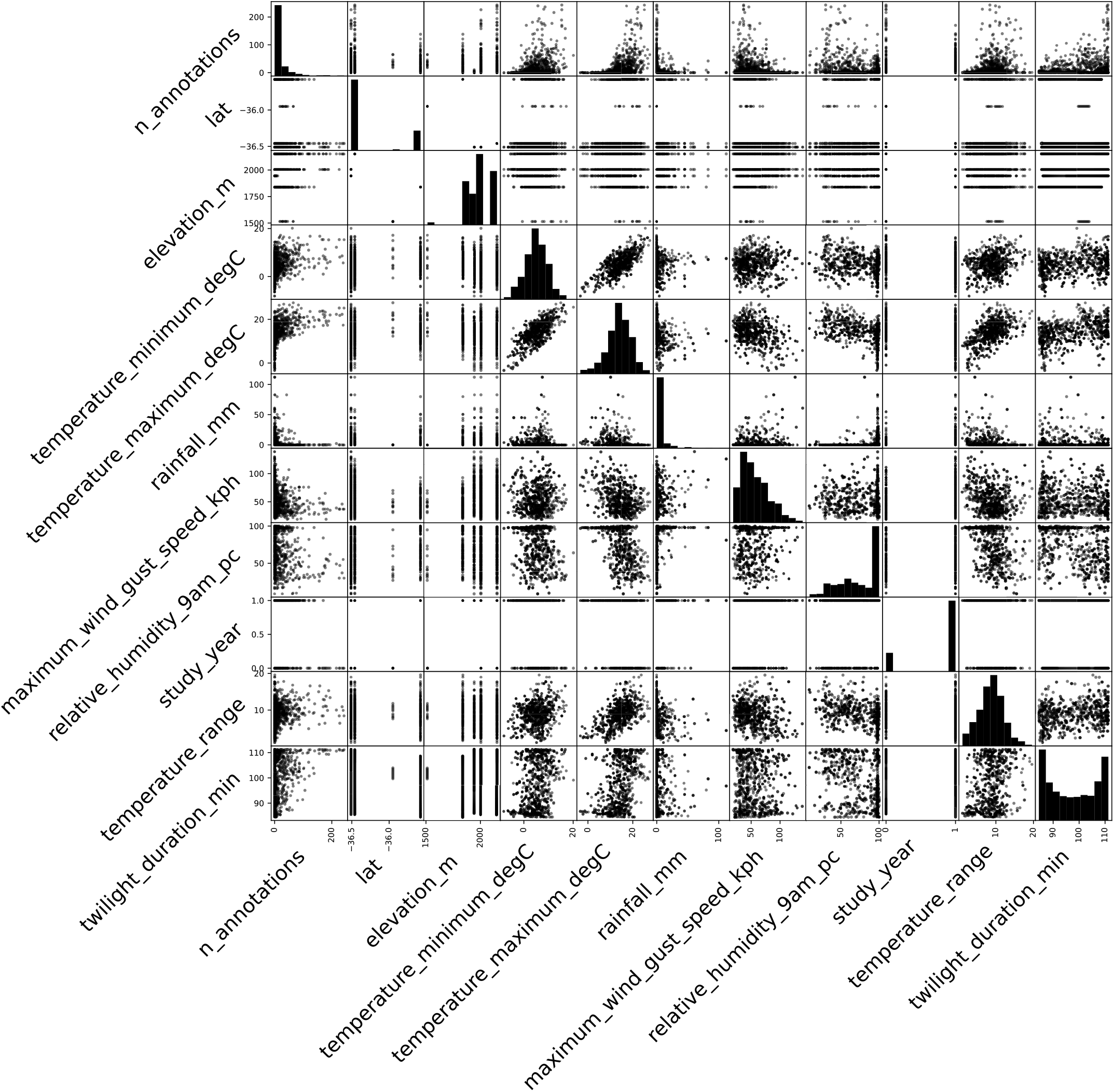
Scatter matrix of Bogong evening twilight flight covariates.

### A.3: Bogong moths flying during bushfire

**Figure A.3.**
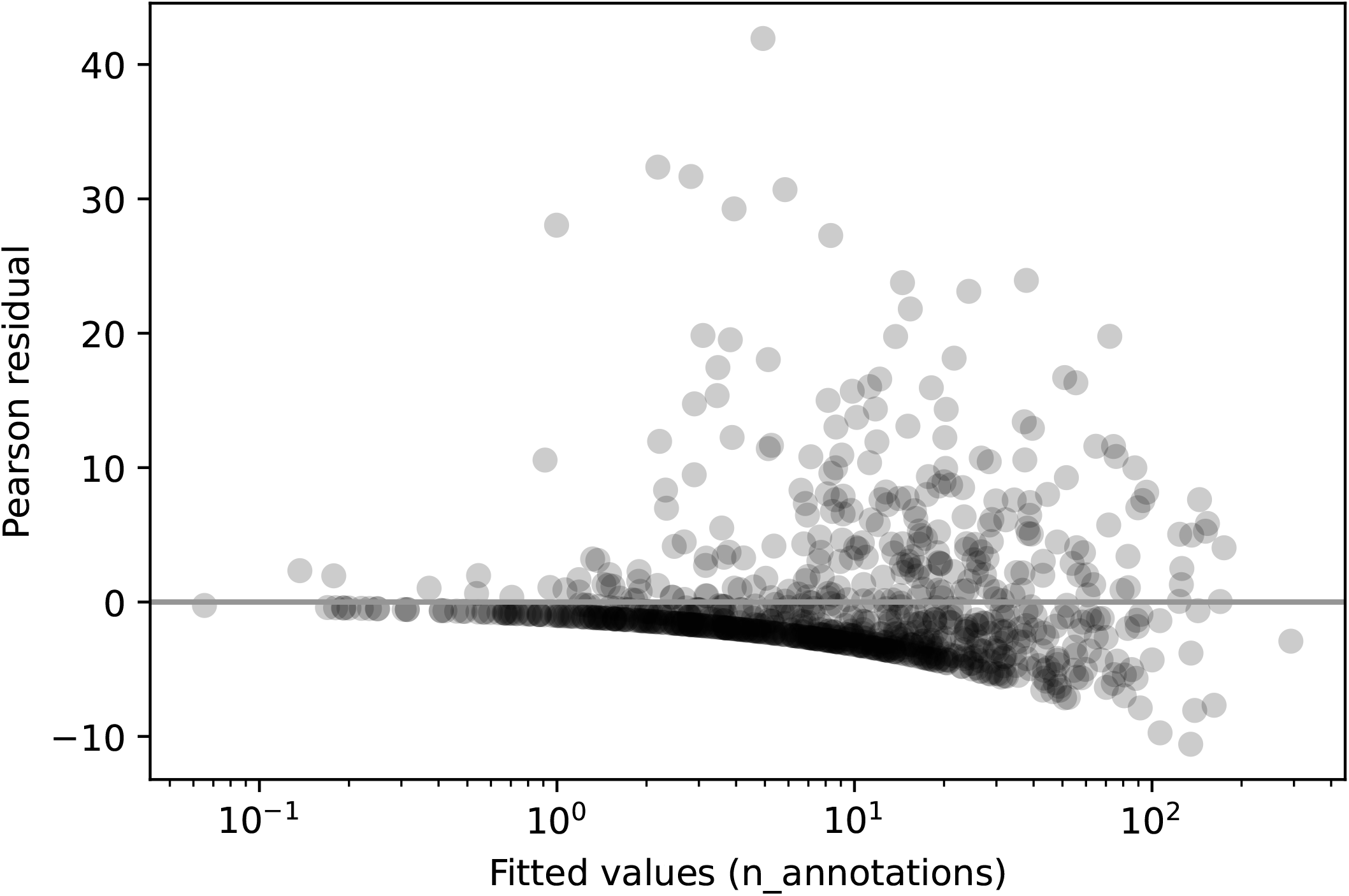
Pearson residuals versus predicted evening twilight detection count for Poisson GLM of detections against (in order of effect size); elevation, maximum daily temperature, day length, maximum wind speed, study year, temperature range, 9 am relative humidity, latitude, minimum temperature, and rainfall.

**Figure A.4.**
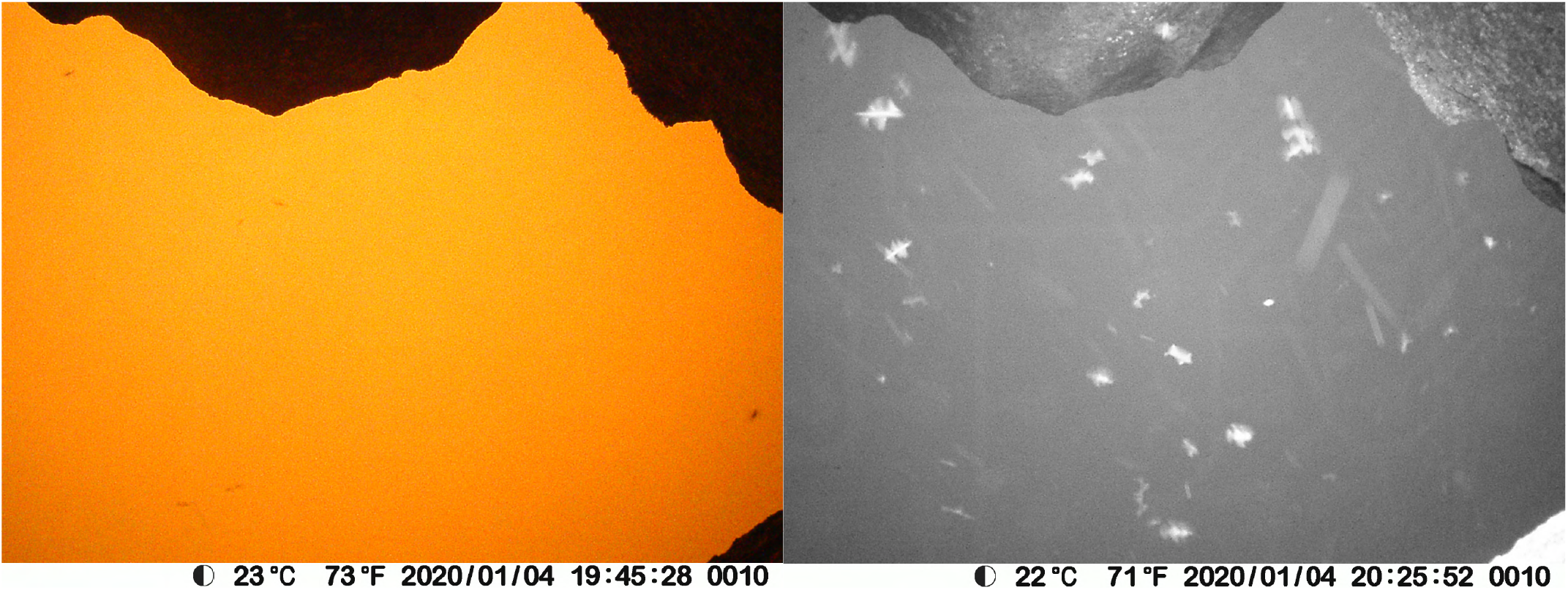
Bogong moths flying during bushfire outside aestivation cave near the top of Ken Green Bogong on 4^th^ January 2020. **Left:** Photograph taken by camera, shortly before switching to “night mode.” The air is thick with smoke, leading to the orange colour. Dark specks in the air are likely Bogong moths. **Right:** Photograph taken by the same camera, once it had switched to “night mode,” with infra-red flash. Flying Bogong moths are clearly visible.

